# Circadian modulation by time-restricted feeding restores brain transcription and slows amyloid deposition in a mouse model of Alzheimer’s disease

**DOI:** 10.1101/2022.10.07.511346

**Authors:** Daniel S. Whittaker, Laila Akhmetova, Haylie Romero, David K. Welsh, Christopher S. Colwell, Paula Desplats

**Affiliations:** Department of Neurosciences, University of California San Diego, La Jolla, CA, USA; Center for Circadian Biology, University of California San Diego, La Jolla, CA, USA; Department of Psychiatry, University of California San Diego, La Jolla, CA, USA; Veterans Affairs San Diego Healthcare System, San Diego, CA, USA; Department of Psychiatry and Biobehavioral Sciences, University of California Los Angeles, Los Angeles, CA, USA; Department of Pathology, University of California San Diego, La Jolla, CA, USA

**Author notes:** Corresponding author, Paula Desplats, Ph.D., University California San Diego, 9500 Gilman Drive, MTF346, MC0624, La Jolla, CA 92093.

**Keywords:** Alzheimer’s disease, circadian rhythms, feed/fast cycle, time-restricted feeding, rhythmic transcription, amyloid plaque deposition, neuroinflammation, APP23, hippocampus transcriptome, Alzheimer’s disease-associated genes

## Abstract

Alzheimer’s disease (AD) is a tragic neurodegenerative disease affecting more than 5 million Americans. Circadian disruptions impact nearly all AD patients, with reversal of sleep/wake cycles and agitation in the evening being common disturbances that manifest early in disease. These alterations support a role for circadian dysfunction as a driver of AD, emphasizing a critical need to investigate the therapeutic potential of circadian-modulating interventions. One of the most powerful regulators of the circadian system is the daily feed/fast cycle. Here we show that time-restricted feeding (TRF) without caloric restriction, improved key disease components including behavior, disease pathology and transcription in the APP23 mouse model of Alzheimer’s disease. We found that TRF had the remarkable capability of simultaneously reducing amyloid deposition, increasing Aβ42 clearance, improving sleep and hyperactivity, and normalizing transcription of circadian, AD and neuroinflammation-associated genes in APP23 mice. Thus, our study unveils for the first time that circadian modulation through timed feeding has far-reaching effects beyond metabolism and affects the brain as the substrate for neurodegeneration. Since the pleiotropic effects of TRF can substantially modify disease trajectory, this intervention has immediate translational value, addressing the crucial need for accessible approaches to reduce or halt AD progression.

## INTRODUCTION

Alzheimer’s disease (AD) is a devastating neurodegenerative disorder affecting the lives of more than 5 million Americans and their families, and it is the biggest forthcoming health challenge in the United States ^1^. Beyond accumulation of beta-amyloid and phosphorylated Tau proteins in the brain, disturbance of circadian rhythmicity is a common complaint for more than 80% of AD patients, evidenced by altered sleep/wake cycles and behaviors, such as increased cognitive impairment and confusion in the evening (a.k.a. sundowning), as well as difficulty in falling and staying asleep ^2,3^. These factors are the leading cause of nursing home placement and are also associated with decreased survival ^4^. Treatments delaying AD progression remain elusive; thus, approaches that prolong patient independence and daily functioning are likely to have a great impact in clinical care.

Emerging evidence, including studies from our group, suggests that circadian alterations occur earlier in disease progression than previously estimated and may directly aggravate pathology ^5,6^. Circadian rest-activity patterns are fragmented in preclinical stages of AD, and weakened circadian activity patterns increase the risk of dementia and precede cognitive impairment by several years ^7,8^. However, the pathways that mediate circadian dysregulation in AD and how they are modulated are not yet fully defined.

Circadian (ca 24 h) clocks coordinate the daily temporal organization of physiology and behavior through tightly regulated transcriptional programs. In addition, clock transcription factors can modulate downstream targets outside the core clock, thus imposing rhythmicity in up to 50% of the transcriptome ^9^. Cell-autonomous clocks reside in various brain regions, including those severely affected by AD like hippocampus and frontal cortex ^10,11^. Clock misalignment is associated with poor health and AD risk factors including diabetes, cardiovascular diseases, inflammation, and sleep disorders ^12^. Deletion of core circadian clock genes *Bmal1* and *Per1* in mouse brain triggers synaptic degeneration, impaired cortical connectivity, oxidative damage, behavioral abnormalities and memory impairment, highlighting the impact of circadian alterations on cognitive function and neuronal viability ^13,14^.

Modulation of the circadian clock as a strategy to improve health outcomes is gaining momentum. Particularly, the daily feed/fast cycle provides a strong stimulus for the synchronization of metabolic and behavioral functions even in the absence of a functional central circadian clock ^15–18^. We recently demonstrated that time-restricted feeding (TRF; 6 h feed, 18 h fast cycles) can improve sleep/wake cycles, motor performance, and inflammation in mouse models of Huntington’s disease ^19,20^. Furthermore, human studies applying analogous “Time Restricted Eating” paradigms showed changes in diurnal patterns of cortisol, and in the expression of circadian clock genes and autophagy markers in blood ^21^.

The mechanisms through which TRF conveys the observed benefits have not been fully elucidated and are likely pleiotropic. The alterations brought about by TRF may result from changes in metabolite interactions, bioenergetic pathway responses, modulation of circadian rhythm timing and strength, epigenetic modifications, and effects on food-anticipatory activity and reward circuits ^22–28^.

In this study, we identified progressive circadian disruptions in the APP23 transgenic mouse model of AD, including excessive wakefulness, altered behavioral circadian rhythms and hyperactivity and severe deregulation of the oscillatory expression patterns of many genes associated with AD pathology and neuroinflammation in the hippocampus. We evaluated whether modulation of the circadian clock at early-disease stages mitigates behavioral and transcriptional alterations and ameliorates pathology. We report that a TRF protocol limiting food access to 6 h -aligned to the active phase-improved diurnal locomotor activity patterns and behavioral circadian rhythms and increased total sleep, without caloric restriction. Importantly, we determined that TRF normalized the transcription of genes associated with AD, neuroinflammation, lipid processing, and autophagy, as well as improving the rhythmicity of core clock genes in the hippocampus of APP23 TG mice. Critically, TRF had a major impact on neuropathology, with APP23 TG treated mice displaying a significant reduction in plaque burden, amyloid deposition, and a reversal of circulating biomarkers of AD. Overall, our results demonstrate that application of TRF can forestall behavioral and molecular disruptions that aggravate AD pathology in mouse models.

## RESULTS

### AD mice present disruption in sleep, activity and circadian regulation

APP23 TG mice develop progressive pathology starting with sparse amyloid plaques in the cortex and hippocampus around six months of age ^29^. Importantly, TG mice at this age already display increased fragmentation of sleep during the light phase and some cognitive alterations ^30^. Sleep disruptions continue to worsen with age and pathology, and we observed that 11-month-old APP23 TG mice showed decreased mean total sleep and significant hyposomnia during the active phase compared with NTG controls (Figure 1A-B; Ext. data Fig.1a-b) and altered activity rhythms with excessive cage activity during the dark phase (Figure 1C-D; Ext. data Fig.1c-d).

**FIGURE 1.**
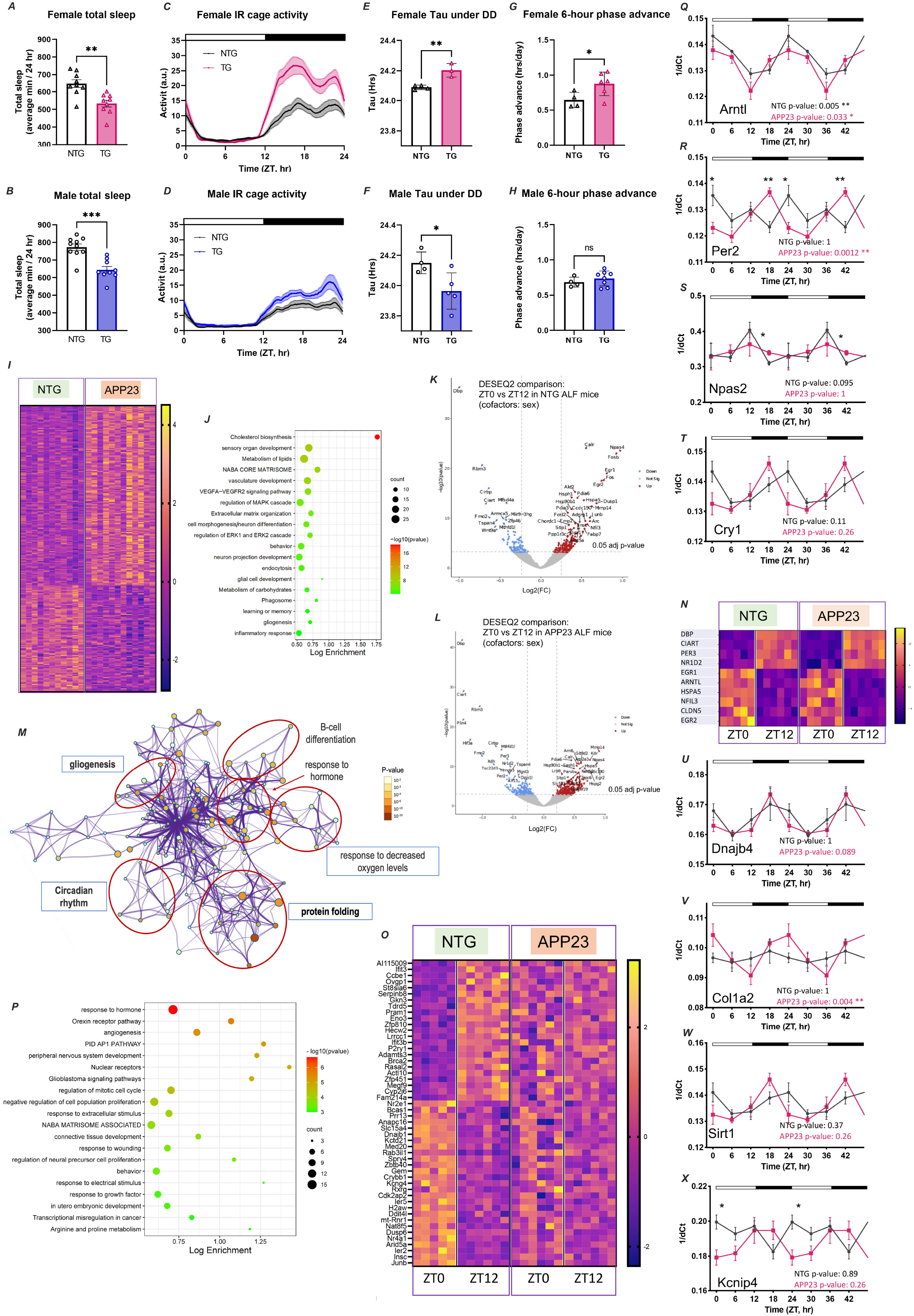
AD mice show alterations in sleep, activity, circadian rhythms and brain transcription. **A-B**. Total sleep is represented as average minutes per 24 h from data collected over two 24 h sleep-wake cycles. **C-D**. Activity recordings reported in 3-min bins averaged from 7-10 days of activity. **E-F**. Tau (circadian period length) under DD was calculated from activity onset times and is presented in hour units. **G-H**. Phase shift was calculated as [(Δh “activity onset” to “entrained onset”)/days to entrainment] and is presented as the shift in h/day. Top bar in waveforms represents the light-dark cycle. Bar graphs represent individual data plots with standard error of the mean. Statistical significance represents the comparison between TG and NTG mice as per unpaired Students’ *t*-test *p≤0.05; **p≤0.01; ***p≤0.001. **I**. Heatmap of significant DEGs in APP23 TG vs NTG mice based on RNA-Seq analysis of hippocampus tissue using DESEQ2; threshold adj.p≤0.2. Each row is one gene and expression is represented by z-score. **J**. Gene ontology terms for genes differentially expressed in TG mice using Metascape. **K-L**. Transcriptional profiling by RNA-Seq shows diurnal rhythms of expression in both APP23 TG (K; n=12) and NTG mice (L; n=13) hippocampus. Depicted as volcano plots at adj.p≤0.05; red denotes increased and blue decreased at ZT0 in comparison to ZT12. **M**. Pathway enrichment analysis of rhythmic genes using Metascape and showing enrichment in functions associated with neurodegeneration and circadian clock. **N**. Expression of circadian clock genes in NTG and TG mice, presented as heatmap sorted by phase of gene expression in z-scores. **O**. Rhythmic expression of some genes is obliterated in APP23 TG mouse hippocampus, as shown by heatmap sorted by phase of expression in NTG and represented by z-scores. **P**. GO analysis of the genes that lost rhythmicity in TG mice, showing top enriched functions in Metascape, including pathways associated with transcription regulation, sleep, and behavior. **Q-X**. Alterations in the rhythmic patterns of expression of core clock (Q-T) and AD-associated genes (U-X) were detected in APP23 mice by qPCR using samples taken every 6 hr. Transcript abundance at each time point is expressed as inverse delta Ct. The periodicity p-value is denoted on each plot for NTG and TG. Values are double plotted. Statistical significance represents the comparison between TG and NTG mice at a single timepoint as per unpaired Students’ *t*-test, *p≤0.05; **p≤0.01.

Evaluation of endogenous circadian activity rhythms in constant darkness conditions (DD) showed alterations in circadian period, with TG females showing longer and males exhibiting shorter periods than NTG controls (Figure 1E-F; Ext. data Fig.2i-l). TG mice also showed increased fragmentation of activity under DD, indicated by reduced activity bout lengths and a trend towards increased activity bout number, with no changes in total activity in LD vs DD (Ext. data Fig.2a-f). Furthermore, female TG mice re-entrained significantly faster than NTG controls in response to a 6 h phase advance (a behavior not observed in males; Figure 1G-H; Ext. data Fig.2m-p), and both female and male TG mice showed significant activity suppression in response to negative light masking, similar to NTG animals (Ext. data Fig.2g-h). Taken together, these observations demonstrate that APP23 TG mice present sleep and circadian disruptions (without exhibiting impairments in their response to light) at early disease stages that precede substantial amyloid pathology.

### AD mice show dysregulation of gene expression in the hippocampus

To investigate early transcriptional alterations associated with AD pathology in the hippocampus of APP23 TG mice, we applied RNAseq and performed differential gene expression analysis in comparison to NTG animals. We identified 258 differentially expressed genes (DEGs) in TG mice, including 28 AD-genes overlapping with AMP-AD and DisGENET Databases (Figure 1I-J; Ext. data Table 1). Gene ontology (GO) analysis revealed involvement of DEGs in cholesterol biosynthesis, core matrisome and extracellular matrix (ECM) organization, vasculature development, neuron and glial cell functions, memory, inflammation, endocytosis and phagocytosis; with significant enrichment for astrocyte markers (Ext. data Table 2). Upstream regulator analysis predicted activation of MAP2K5 kinase (z-score=4.2; p-value of the overlap=1.68E-30) and SREBF2 (z-score=3.8; p-value of the overlap=1.58E-24), both implicated in cholesterol synthesis and insulin metabolism. Among the top inhibited regulators, we identified the group of β-adrenergic receptors (ADRB; z-score=−3.3; p-value of the overlap=6.12E-9), which mediate norepinephrine signaling facilitating synaptic plasticity and memory formation and that have been extensively investigated as therapeutic targets for AD ^31^.

### Impaired daily rhythms of transcription in AD-mice hippocampus

To evaluate potential alterations in rhythmic transcription in the hippocampus of TG mice, we analyzed RNAseq data from brain samples obtained at ZT0 and ZT12 (n=3/4 mice per time point per genotype). We identified 248 genes in NTG mice and 623 genes in TG mice that were differentially expressed between ZT0 and ZT12 (Figure 1K-L; Ext. data Table 3). Pathway analysis of 127 DEGs that overlapped between NTG and TG mice showed enrichment in protein folding, response to hormones and growth factors, and circadian rhythms (Figure 1M). The latter included the core clock genes *Dbp, Nr1d2, Per3*, and *Arntl*, thus validating our approach to identify rhythmic genes (Figure 1N). Markedly, we identified 121 genes that displayed time-of-day-specific expression in NTG mice but were arrhythmic in TG animals (Figure 1O; Ext. data Table 3). Pathway analysis indicated that genes that lost diurnal variation in TG mice were involved in response to hormones, angiogenesis, ECM, and orexin signaling, and showed enrichment for oligodendrocyte cell markers, thus offering novel insights into the pathways where circadian dysregulation impacts AD pathology (Figure 1P). These results demonstrate that a large fraction of genes perturbed in AD exhibit time-of-day-specific expression, underscoring the impact that broken circadian rhythms could have in disrupting the hippocampal transcriptome.

After finding that transcriptional rhythms were perturbed in TG mouse hippocampi, we further examined core clock gene regulation by collecting additional tissue at ZT0, ZT6, ZT12 and ZT18 and applying qPCR followed by JTK_Cycle analysis. We detected alterations in the expression of core clock genes in TG mice, including reduced amplitude of *Npas2*, a phase advance of *Cry1* and *Per2*, which also showed increased rhythmicity, and a trend towards overall decreased expression of *Arntl* (Figure 1Q-T; Ext. data Table 4).

Using this extended time point sample set, we also tested the rhythmic expression of selected AD-associated genes that ranked as top DEGs in the RNAseq analysis. We observed a phase advance in the acrophase of *Kcnip4* and *Sirt1*, and increased amplitude of *Col1a2* and *Sirt1*, with *Col1a2* and *Dnajb4* showing significant strengthening of rhythmicity in TG mice (Figure 1U-X; Ext. data Table 4). These results demonstrate that the hippocampal circadian clock is altered early in AD progression in the APP23 mouse model, potentially triggering downstream dysregulation of rhythmic genes, many of which are directly associated with pathology.

### TRF effectively modulates metabolic markers and ketone-responsive genes in AD mice

Having detected disruptions in activity rhythms, sleep, and brain transcription that potentially contribute to pathology in APP23 TG mice, we evaluated whether a circadian and metabolic intervention could ameliorate AD phenotypes. TG and NTG littermate mice were randomly assigned to either *ad libitum* feeding (ALF) or time-restricted access to food (TRF) with a 6 h feeding/18 h fasting regimen, and the feeding window aligned to the middle of the active period (ZT15-ZT21; Figure 2A).

**FIGURE 2.**
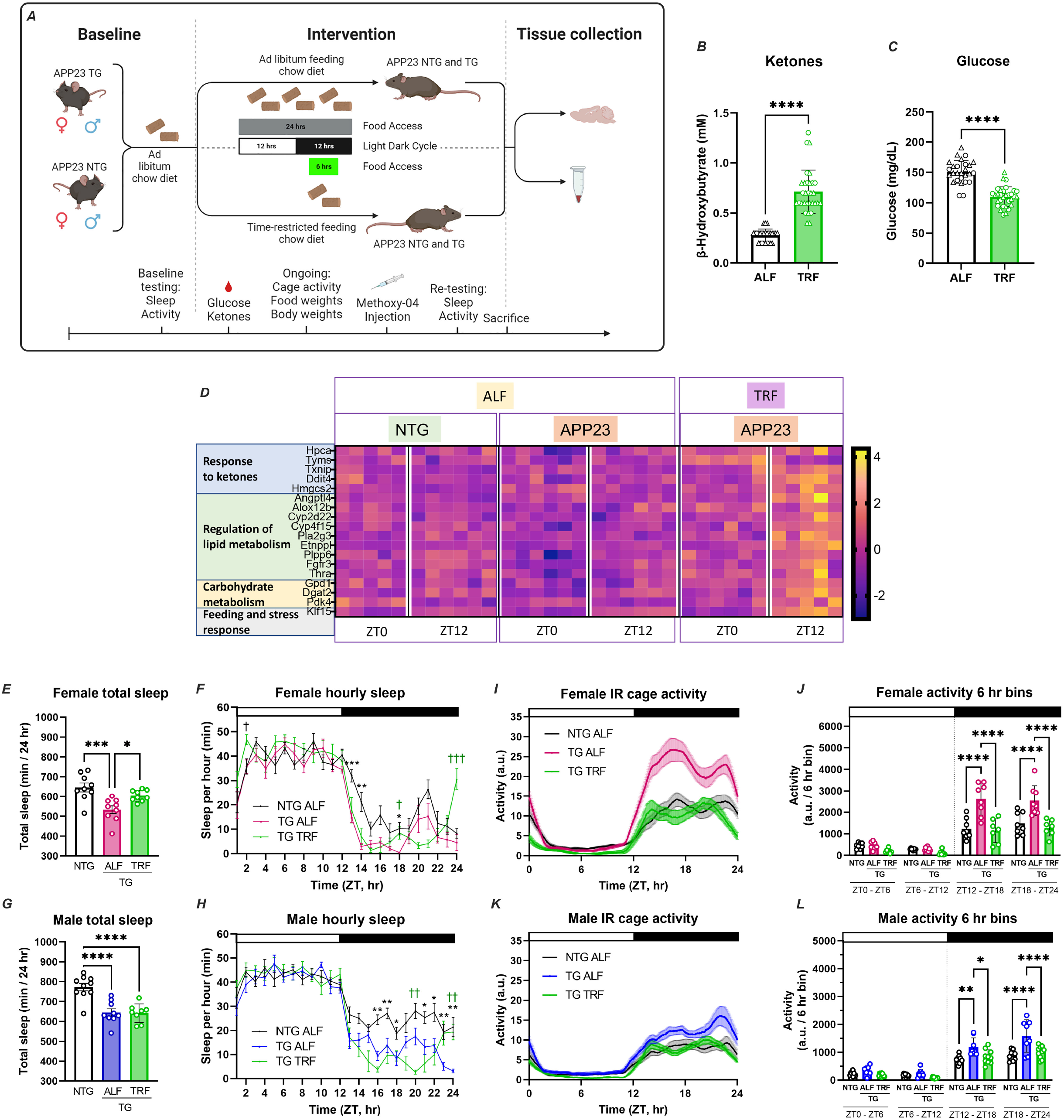
Time-restricted feeding induces metabolic changes, modulates brain transcription and rescues behavior and sleep in AD mice. **A**. Schematic representation of TRF intervention indicating feeding models and assay/evaluations performed. **B-C**. Significant changes in β-hydroxybutyrate and glucose in TRF treated animals (n=31) in comparison to ALF mice (n=27), as detected in blood and presented as individual values with standard error of the mean. Statistical significance as per unpaired Students’ t-test. ◯ female; △ male. **D**. The expression of a set of metabolism-associated genes was modulated by TRF in APP23 TG mice in a phase-dependent fashion, as shown by heatmap based on z-score of normalized reads and sorted by phase of expression in NTG mice. **E-H**. Total sleep is represented as average minutes per 24 h and sleep as average sleep per 1 h bin. Data was analyzed averaged over two 24 h sleep-wake cycles. Statistical significance represents one-way ANOVA with Tukey’s multiple comparisons test (E and G), or two-way ANOVA with Tukey’s multiple comparisons test (F and H) **I-L**. Activity recordings reported in 3-min and 6 h bins averaged from 7-10 days of activity. Statistical significance represents one-way ANOVA followed by Šídák’s multiple comparisons test (J and L). Top bar in waveforms represents the light-dark cycle. Bar graphs represent individual data plots with standard error of the mean. *p≤0.05; **p≤0.01; *** p≤0.001; **** p≤0.0001.

The metabolic effects of TRF are well described, including modulation of glucose and ketone pathways ^21,32,33^. In the present intervention, mice under TRF showed significantly increased β-Hydroxybutyrate and reduced glucose levels in blood at ZT14 in comparison to mice under ALF conditions (Figure 2B-C). Importantly, mice in all groups consumed equivalent volumes of food and showed no significant differences in body weight (Ext. data Fig.3a-d), establishing that any observed effects were not mediated by caloric restriction. In addition, ketone-response genes and transcripts associated with lipid and carbohydrate metabolism were differentially expressed in the brains of TG mice under TRF, as shown by RNA-Seq data. Importantly, TRF imposed time-of-day-specific changes in the expression of these genes, with the largest variations occurring at ZT12, after 15 h of fasting. Among these genes, we detected the transcription factor *Klf15*, which binds hippocampal glucocorticoid receptors and whose diurnal rhythmicity is known to be directly regulated by feeding ^34^ (Figure 2D). These findings confirm that TRF was effective in modulating the appropriate blood metabolites, and, importantly, that these peripheral changes were able to induce specific and temporal transcriptional responses in the brain. Moreover, these data demonstrate that despite ongoing brain pathology, APP23 TG mice can respond to this TRF intervention.

### TRF improves sleep and restores activity rhythms in AD mice

We next evaluated the impact of TRF on behavior in APP23 mice. Temporal restriction of food intake for three months improved different aspects of sleep in females vs. males. Female TG mice on TRF showed increased total sleep, reaching levels like those observed in NTG animals (Figure 2E-F). While no changes in total sleep were observed in TG males under TRF, these animals still showed increased mid-day wakefulness (Figure 2G-H). Importantly, both TG females and males under TRF showed improved sleep at ZT24, the beginning of the sleep phase (Figure 2F and H).

Markedly, TRF rescued specific behavioral abnormalities in both female and male APP23 TG mice, which no longer exhibited active phase hyperactivity, and whose activity patterns became indistinguishable from those observed in NTG mice (Figure 2I-L).

### TRF modulates hippocampal transcription and pathways related to AD and inflammation

TRF effects have been extensively investigated in the liver, a metabolic hub with strong circadian rhythmicity. However, the potential effects of TRF in the brain are unknown. To this aim, we first performed differential gene expression analysis of TRF vs. ALF mice, combining APP23 TG and NTG mice for increased power. Our analysis revealed 499 DEGs that respond to TRF in the hippocampus, which presented the same direction of expression changes across genotypes. GO analysis showed these DEGs were significantly enriched for myelination, lipid metabolism, microtubule organization, and blood vessel morphogenesis (Ext. data Fig.4; Ext. data Table 5). Genes upregulated in response to TRF appeared regulated by the androgen receptor (AR; p-value=4E-5) and the transcription factor NFY (p-value=4E-5); whereas NFKB (p-value=2E-4) and XBP1 (p-value=7E-4) were enriched regulators for genes attenuated by TRF. From this group of TRF-modulated genes, 57 showed differential expression between ZT0 and ZT12 in NTG ALF mice; and 78 genes are reported to exhibit circadian rhythmicity in the mouse hippocampus, with 34 of these overlapping ^35^. Thus 22% of genes that respond to TRF are rhythmic (112 out of 499; Ext. data Table 5).

We next focused on identifying TRF-induced changes in gene expression in APP23 TG mice which may specifically impact AD pathology. We used two microarrays from NanoString^®^ focused on AD and neuroinflammation ^36–38^. We determined that 86 AD genes and 100 neuroinflammation genes functionally associated with myelination, transmitter synthesis and storage, autophagy, cytokine remodeling, and adaptive immune response were affected by TRF treatment (Figure 3A-J; Ext. data Table 6). Furthermore, unbiased transcriptome-wide analysis using RNA-seq data revealed 152 genes responding to TRF in TG mice in cross-sectional analysis; while an additional 85 genes changed only at ZT0 and 28 genes changed at ZT12, thus totaling 265 unique genes modulated by TRF in the hippocampus of TG mice (Figure 3K-M; Ext. data Table 7). Notably, 44 of these genes have been previously associated with AD. Functionally, these 265 genes were enriched in pathways that converge into AD pathology, including ECM and core matrisome, vasculature organization, glycosylation, gliogenesis, PI3K-Akt-mTOR signaling, Aβ clearance, myelination, and ketone responses (Figure 3N; Ext. data Table 8). Additionally, the genes modulated by TRF clustered with oligodendrocyte (p-value 1.4E-17) and vascular smooth muscle cell markers (p-value 4.2E-9), suggesting this intervention could impact myelination and the integrity of the blood-brain barrier. Analysis of upstream factors that may co-regulate the expression of TRF-responsive genes in TG mice pointed to PPARA (p-value=3.2E-5) and RELA (p-value=3.2E-5), which directly interact with core clock genes ^39,40^. PPARA is activated in response to fasting and integrates energy metabolism with the clock. Lastly, gene set enrichment analysis (GSEA) further implicated HMGB1, a multifunction transcription factor that regulates chromatin integrity, autophagy, and inflammation ^41^, in the modulation of 36 TRF-responsive genes (13.6% of total genes in set; FDR qvalue=2.47E-8); whereas another set of 33 DEGs (12.5 %) were enriched for PAX4 (FDR qvalue=2.47E-08), a transcription factor also implicated in circadian regulation ^42^.

**FIGURE 3.**
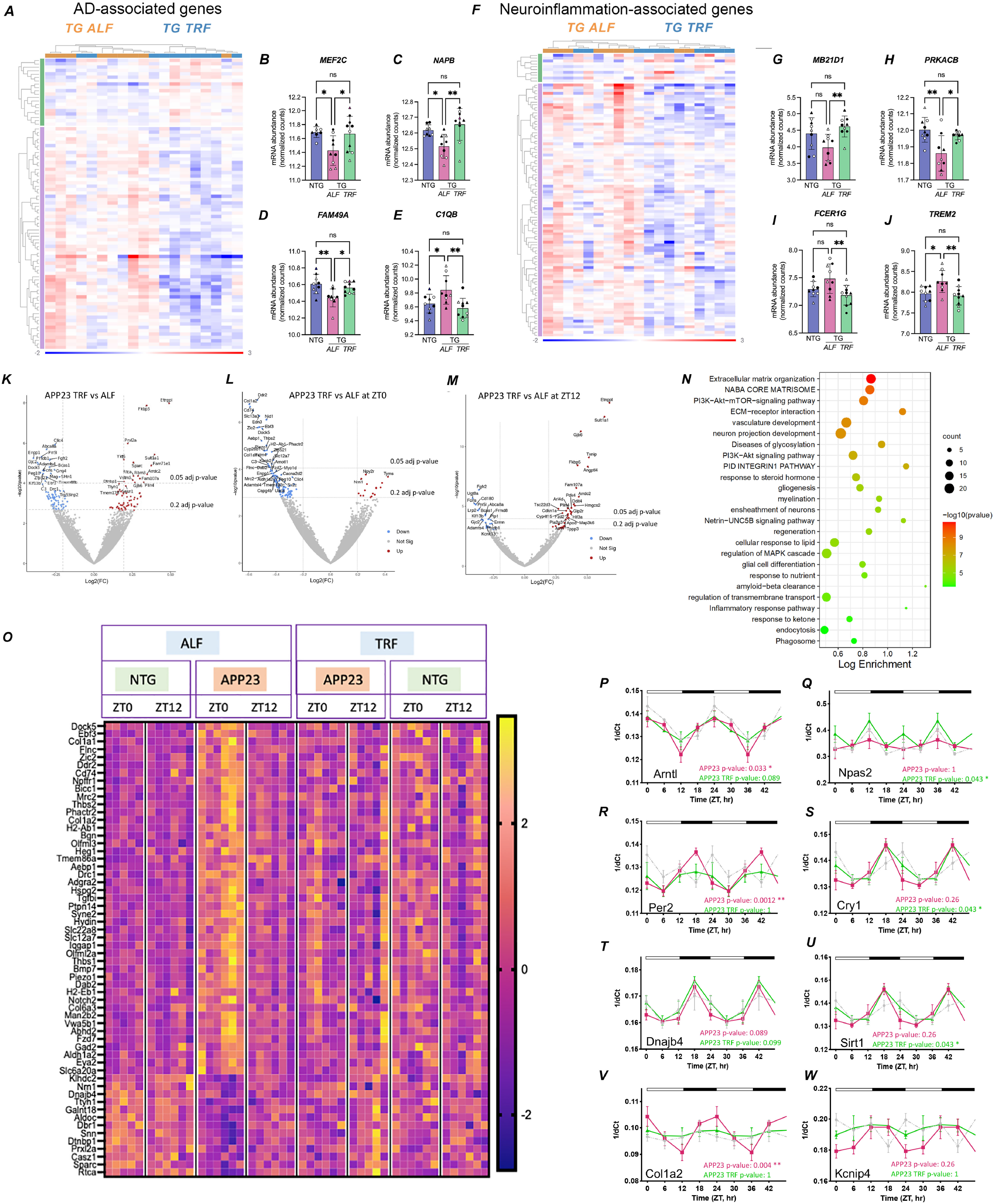
Effects of TRF on hippocampus transcription and diurnal rhythms of gene expression. **A-J**. Differential expression of genes associated with Alzheimer’s disease (A-E) and neuroinflammation (F-J) in APP123 TG mice under TRF (n=10) vs ALF regimen (n=9) and as detected by NanoString^®^ panels. Heatmaps sorted by fold-change after normalization and analysis in Rosalind. Comparison based on treatment, controlling by genotype, sex, and time of collection. Examples of gene expression are shown by box plots for the top changed genes in each group (B-E and G-J) and are based on normalized counts and presented as individual values with standard error of the mean. Statistical significance as per unpaired Students’ t-test *p≤0.05; **p≤0.01. ◯ female; △ male; white symbols for samples taken at ZT0; black symbols for samples taken at ZT12. **K-M**. Significant DEGs based on RNA-Seq analysis of hippocampus tissue using DESEQ2; threshold adj.p≤0.2; shown in volcano plots. Red denotes increased and blue decreased expression in APP23 TG TRF (n=12) in comparison to ALF mice (n=12). **N**. Pathway enrichment analysis using Metascape, representing top significantly enriched functions for genes modulated by TRF in TG mice. **O**. TRF partially rescued transcriptional alterations observed in APP23 TG mice as shown in heatmap of 58 genes sorted by significance of the differential gene expression in APP23 TG mice versus NTG animals. **P-W**. TRF also improved the circadian pattern of expression of core clock genes in APP23 TG mice (P-S) and AD-associated genes (T-W), detected by qPCR using samples taken every 6 hr. Transcript abundance at each time point is expressed as inverse delta Ct. Periodicity p-value is denoted on each plot for TG ALF and TG TRF. Values are double plotted. Statistical significance represents the comparison between TG and NTG mice at a single timepoint as per unpaired Students’ *t*-test, *p≤0.05; **p≤0.01.

Importantly, more than 20% (58 genes) of the 265 genes modulated by TRF were aberrantly expressed in untreated APP23 TG mice and were restored by treatment, with their expression levels approaching those observed in NTG ALF animals. This group included AD-associated genes *Col1a2, Cd74, Nrn1* and *Iqgap1* (Figure 3O; Ext. data Table 9). Pathway analysis of genes restored by TRF revealed enrichment for inflammatory response, phagosome activity, integrins, and ECM-receptor interaction. As a result of all these changes, APP23 TG mice subjected to TRF became transcriptionally similar to NTG TRF animals, with only 26 genes showing differences in expression between genotypes, in marked contrast to the 258 DEGs originally identified for the TG vs NTG comparison under the ALF conditions (Ext. data Figure 5; Ext. data Table 10).

Finally, we evaluated whether TRF also ameliorated the alterations in core clock gene expression observed in untreated TG mice. Analysis of the 24 h time point samples by qPCR showed partial reversal of the decrease in *Arntl* expression, restoration of rhythmicity in *Npas2* and improved phasing of *Cry1* (Figure 3P-S). Similarly, TRF modulated rhythmic AD genes, strengthening the rhythmicity of *Sirt1*, and reducing the amplitude of *Kcnip4* and *Col1a2* (Figure 3T-W).

Altogether, these findings indicate that TRF has a profound impact on the brain transcriptome in APP23 TG mice, normalizing patterns of gene expression in the hippocampus. Moreover, the identification of a substantial proportion of TRF-responsive genes that show rhythmic expression, and the role of their predicted upstream regulators AR, NFY, NFKβ, PPARA and PAX4 in modulating core clock genes ^39,42–45^, supports the hypothesis that the pleiotropic effects of TRF in the brain are mediated by a complex cross-talk between the circadian clock and homeostatic systems.

### TRF reduces AD pathology in APP23 TG mice

In view of the effects of TRF in modulating pathways strongly associated with AD pathology we finally evaluated the impact of this intervention on disease trajectory. TRF-treated mice showed sparse, small core (dense) plaques, and intracellular accumulation of amyloid-β (Figure 4A-H), which may represent a pool for the formation of extracellular aggregates later in disease ^46^. Quantification of amyloid aggregates in APP23 TG mice showed that the total area occupied by plaques, and the number of plaques per mouse were both significantly reduced under TRF (Figure 4I-J). Plaque-size analysis showed reduced numbers of plaques of any size under TRF, with more pronounced and significant changes observed for smaller plaques (under 3,000 μm^2^), and potentially indicating changes in the rate of amyloid deposition (Figure 4K). Thus, amyloid pathology in TRF mice appeared to be at earlier stages in comparison to ALF TG mice.

**FIGURE 4.**
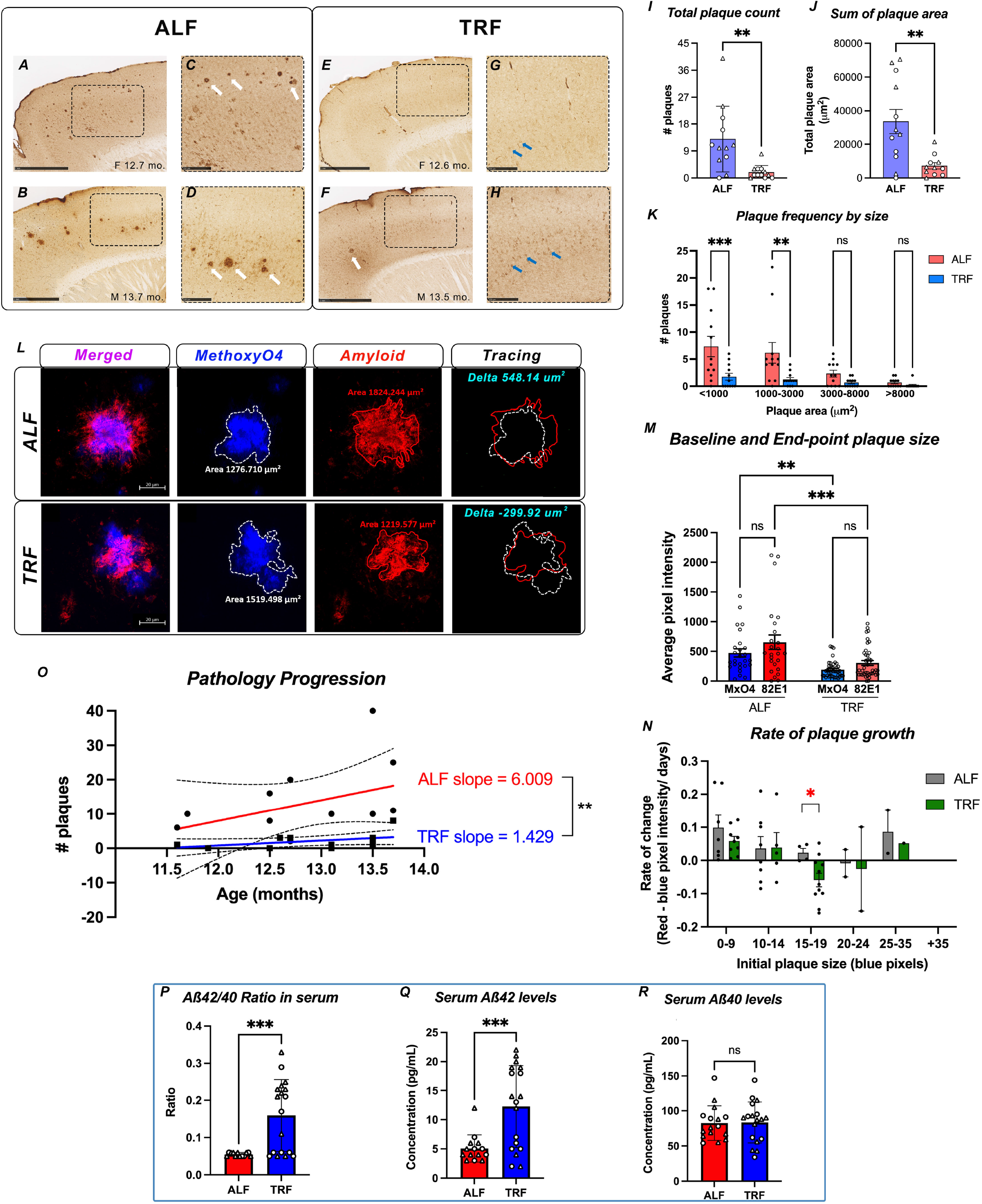
TRF significantly ameliorates amyloid pathology and disease progression in AD mice. **A-H**. APP23 TG mice show progressive amyloid pathology, with plaque accumulation in the frontal and medial cortex. Pathology is substantially reduced for APP23 TG mice in TRF, as sex- and age-matched animals show only sparse plaques (white arrows) and intracellular accumulation of amyloid-β (blue arrows). **C, D, G and H**. Higher magnification of sections indicated in A, B, E and F. **I-J**. Quantification of amyloid plaque counts (I), area occupied by plaques (J), and comparison of plaque number after binning by size (K) in APP23 TG mice in TRF (n=11) vs ALF (n=12). ◯ female; △ male. **L-M**. In vivo analysis of plaque growth in APP23 TG mice. Baseline plaques were labeled by ip injection of Methoxy-X04 (blue signal). End point plaques were detected by immunostaining of sagittal sections with anti-amyloid antibody 82E1 (red signal). Orthogonal projections of z-stacks obtained by confocal microscopy (80x) were used to trace and calculate the plaque area using ZEN Blue digital imaging software. Panels show representative images from single plaques analyzed in one TRF and one ALF TG (L; scale bar = 20 μm) and (M) shows quantification as average pixel intensity per channel from plaques analyzed in 12 mice/ condition. **N-O**. TRF slows down disease progression as seen by rate of plaque growth calculated as the subtraction of the area covered by 82E1 minus area covered by Methoxy-X04 and plotted as bins based on baseline plaque size (N) and comparing the slopes obtained by simple linear regression of plotting total plaque number per mouse as a function of age in APP23 TG mice in TRF and ALF (O). **P-R**. TRF modulates clinically relevant blood biomarkers in treated mice. Total serum levels of amyloid-β fragments 40 and 42 were quantified by ELISA (Mesoscale) in APP23 TG mice in TRF (n=18) vs ALF (n=12) and presented individually or as ratio. ◯ female; △ male. Bar graphs represent individual data plots with standard error of the mean. Statistical significance represents the comparison between TG TRF vs ALF mice as per unpaired Students’ t-test (I-J; O and P-R) or one-way ANOVA with Tukey’s multiple comparisons test (K, M and N)* p≤0.05; **p≤0.01; *** p≤0.001.

To understand whether the dynamics of amyloid-β deposition/clearance were altered by TRF, we labeled the plaques present at baseline via i.p. injection of Methoxy-X04, as reported previously ^47^. Post-mortem immunohistochemical detection of amyloid (using the anti-amyloid 82E1 antibody) then enabled us to quantify longitudinal changes in the number and size of plaques. Orthogonal projections of z-stacked images from Methoxy-X04-labelled plaques (blue channel) and 82E1+ plaques (red channel) were used to determine plaque growth (Figure 4L). Amyloid plaques show a trend towards increased size in TG animals in ALF, which did not reach statistical significance likely due to low levels of initial pathology and over the short treatment period (3-month). In striking contrast, we observed that end-point plaque size was significantly diminished in TRF-treated mice, indicative of TRF impeding the growth of plaques. We also observed a significant reduction in the size of Methoxy-X04-labelled plaques after 3 months on TRF, suggesting that TRF induced clearance of pre-existing plaques (Figure 4M). Overall, plaque growth rate was lower under TRF, especially for medium-sized plaques (Figure 4N). Linear regression analysis showed a positive correlation between plaque load and age for APP23 TG mice in all groups, but, strikingly, there was a significant difference in the elevations of the curves (p=0.0021), with TG mice in TRF showing a 42.8% reduction in the slope (1.429) in comparison to the ALF group (6.009), - demonstrating that overall disease progression was significantly attenuated by treatment (Figure 4O).

Immunostaining and quantification of NeuN+ nuclei revealed significant, albeit modest, neuronal loss in the hippocampus of TG animals, with a trend towards reduced neuronal counts in the cortex. These deficiencies were partially restored by TRF (Ext. data Figure 6a-e). Similarly, a trend towards increased Iba1+ microglia abundance was apparent in APP23 TG in both cortex and hippocampus, although TRF treatment did not significantly affect microglial cell number (Ext. data Figure 6f-j). Further, we observed increased astrogliosis in the cortex of TG mice, which was also reduced by TRF treatment, whereas no significant changes were detected in the hippocampus (Ext. data Figure 6k-o).

Finally, we explored whether the observed effects of TRF on brain pathology were reflected by circulating biomarkers in APP23 TG mice, by measuring Aβ40 and Aβ42 levels in serum using an anti-human amyloid 6E10 multispot assay. Strikingly, we discovered that the Aβ42/40 ratio was significantly elevated by TRF in TG mice. This change was driven by higher Aβ42 concentrations in the TRF-treated animals and indicates increased Aβ clearance, thus providing further insights into the mechanisms by which TRF significantly reduces pathology in APP23 TG mice (Figure 4P-R).

## DISCUSSION

Disruptions in sleep and daily activity patterns are highly prevalent among AD patients, and emerging data indicate that these changes occur early in disease progression, potentially representing causal factors rather than a consequence of neurodegeneration.

Here, we demonstrate the efficacy of a circadian intervention based on time-restricting feeding in rescuing pathology in the APP23 TG mouse model of AD. We provide ample evidence of the pleiotropic effects of TRF treatment in modulating behavior and sleep, and in normalizing hippocampal gene expression in specific pathways associated with AD and neuroinflammation. Importantly, our results show that TRF can alter disease trajectory by slowing progression of amyloid pathology, as evidenced by reduced plaque load, slower rate of amyloid deposition and increased Aβ42 clearance. While the effects of TRF in synchronizing and strengthening circadian rhythmicity in metabolic hubs like the liver are very well known, our study unveils for the first time that circadian modulation through time-restricted feeding has more far-reaching effects, affecting the brain as the substrate for neurodegeneration.

In addition to increasing total sleep and resolving sundowning-like hyperactivity, TRF specifically improved sleep at the beginning of the daily sleep period. Among the most common sleep disturbances in AD patients are excessive daytime sleepiness, agitated behavior after sundown, and sleep disruptions, including difficulty in falling asleep and difficulty staying asleep (sleep fragmentation), all associated with declining cognitive performance and reduced white matter volumes ^48^. These alterations are viewed as disruptions in the circadian organization of sleep and wakefulness, which have been reported to increase the accumulation of Aβ. At the same time, Aβ accumulation and the resulting neuronal and synaptic damage can drive sleep disturbances and dysfunction of circadian clocks during the preclinical phase of AD, thus fueling a vicious cycle between AD pathology and circadian disruptions ^6^. Here we show that APP23 TG mice lose diurnal oscillation in genes regulated by the orexin signaling pathway, which may partly explain the hyperexcitability we observed in the mice and their reduced sleep. Orexin is a rhythmic neurotransmitter expressed in the hippocampus that modulates sleep, excitability, reward, and motivated behaviors. Notably, orexin and its receptors are diminished in AD patients, contributing to sleep disturbances ^49–51^.

Impairments in the light input pathway have been reported in several AD mouse models ^52,53^, but responses to light were intact in APP23 TG mice in the current study, suggesting that the observed circadian impairments were not a result of light input deficits. In line with TRF acting as a synchronizing stimulus (zeitgeber), we saw rhythmic behavioral and transcriptomic changes in response to TRF. This may have significance for AD patients, where pathology in retina and light input pathways have been reported, and which may contribute to light response impairment and circadian rhythm dysfunction ^54^. Whereas light therapy may help some AD patients with the most severe rest-activity disruptions ^55^, TRF may benefit a larger patient population, representing an efficacious approach to ameliorating circadian rhythm dysfunction.

Entrainment of circadian clocks by timed feeding and arousal are well studied ^56^, yet the benefits of the TRF protocol presented in this study extended beyond transient arousal at the time of feeding in the dark phase. Rather, the impact of TRF was sufficient to drive transcriptional changes in several interconnected pathways, including ECM and vascular remodeling, glycosylation, lipid and cholesterol dynamics, vesicle trafficking, autophagy, protein degradation, Aβ clearance, neuroglial functions, and inflammation, which collectively restored systems disrupted by AD pathology. Further, disrupted diurnal oscillations in a significant number of genes in the TG mice were restored, including in clock and AD-associated genes. Cholesterol pathways were strongly disrupted in APP23 TG mice, representing a primary transcriptional target for the action of TRF. This is particularly relevant to AD, as abnormalities in cholesterol-related gene transcription and pathways have been demonstrated to play a role in altered amyloidogenic processing of APP and disease severity ^57,58^. The extent of TRF-driven changes in hippocampal gene expression, especially as they relate to pathways that impact AD pathogenesis and circadian disruption, may underlie the breadth of benefits observed in this study.

Shedding light on potential mechanisms mediating the effects of TRF, we identified multiple upstream regulators that may drive the transcription of TRF-responsive genes. Looking at DEGs in both NTG and TG mice under TRF, we identified enrichment for AR and NFY, which directly interact with core clock genes ^43,45^. In addition, our analysis predicted attenuation of NFKB1 signaling. NF-κB belongs to a family of redox-sensing molecules, and is elevated in neurons and glia surrounding amyloid plaques, contributing to neuroinflammation, oxidative stress, and apoptosis in AD brains ^59^. Moreover, NF-κB is regulated by the core clock gene *Cry* ^60^ and interacts with other circadian-regulated systems ^40^. Studies in rodents and humans showed that disruptions in sleep caused overexpression of NF-κB, while interventions that improved the quality of sleep in older adults reduced circulating levels of this TF ^44^. Thus, the improved sleep patterns and the partial restoration of core clock gene expression induced by TRF in APP23 TG mice in our study may have attenuated NF-κB activity, contributing to decreased inflammation and reduced amyloid load.

The finding of PPARA as an upstream regulator of genes modulated by TRF specifically in APP23 TG mice also links the beneficial effects of this intervention with improved circadian regulation. Members of the ligand-regulated nuclear receptor family (PPARs) are rhythmically expressed in mouse tissues, with PPARα and PPARγ directly modulating core clock genes *Bmal1* and *Rev-erb*α, linking circadian rhythms and metabolism ^39^. PPAR-α regulates genes associated with glutamate, dopamine, and cholinergic signaling, and modulates the activity of α-secretase and BACE1, thus directly impacting APP processing and degradation. Importantly, PPAR-α is decreased in the brains of AD patients, driving inflammation, oxidative stress, and alterations in lipid metabolism, all pathways that we showed here to be dysregulated in APP23 TG mice ^61^. Interestingly, NFkB, PPARa, and HMGB1 are being actively investigated as therapeutic targets for AD. Our findings showing the potential of TRF to engage these three pivotal factors offer a new promise for non-pharmacological interventions in the treatment of AD.

The peripheral benefits of various feeding regimens have been explored in previous studies, with a recent paper elegantly demonstrating how a circadian-aligned 30% caloric-restricted feeding regimen leads to complex genome-wide reprogramming of circadian gene expression in the liver, including amelioration of age-related changes and protective effects on lifespan ^62^. In numerous preclinical animal models and human studies, restricting feeding to particular time windows without reducing caloric intake attenuated metabolic diseases like obesity, glucose intolerance, dyslipidemia, and age-related decline in cardiac function ^32,63^. Similarly, a fast-mimicking diet was found to promote regeneration, including rejuvenating immune cells, promoting hippocampal neurogenesis, and increasing lifespan in mice, whereas a companion trial in humans improved regenerative markers and reduced disease risk factor biomarkers ^64^. Specifically for AD, recent work has shown that 40% caloric restriction in APP mice reduced amyloid load in the hippocampus ^65^. Intriguingly, a ketogenic diet was shown to reduce Aβ42 and Aβ40 in APP mice ^66^, and recently nutritional ketosis has been shown to improve the function of aquaporin 4 while reducing astrogliosis ^67^. Whereas this body of work shows a variety of benefits from various feeding approaches, the circadian-aligned TRF without caloric restriction in the current study not only normalizes AD pathology-related and core clock gene expression in the hippocampus, but also ameliorates AD-related behaviors and pathology.

Critically, administration of TRF in the APP23 TG mice resulted in a significant reduction in amyloid load, with increased clearance of Aβ42, resulting in higher Aβ42/40 plasma ratios. In sporadic AD, clearance of Aβ42 into cerebrospinal fluid is reduced as early as 10-20 years before the onset of clinical symptoms and shows an inverse correlation with cortical Aβ burden ^68–70^. Hence, our observation has immediate translational value, as these clinical biomarkers could be used to monitor the efficacy of a time-restricted eating regimen in AD patients.

Restriction of the time window of eating has emerged as an important treatment option with whole-system effects. Here we demonstrate that these interventions can also modulate key disease pathways in the brain. Due to the failure of drug-based strategies targeting amyloid, there is an urgent need for novel approaches to reduce or halt disease progression. Our findings support future studies exploring the therapeutic potential of time-restricted eating as a powerful circadian modulator that could significantly modify disease trajectory in AD patients and which is eminently feasible for integration into clinical care.

## ACKNOWLEDGEMENTS

This study was supported by NIA grant #AG061831 to PD and DKW and by a training fellowship to DSW from NIA grant #5T32AG066596-02. The UCSD Microscopy Core is supported by NINDS grant #P30NS047101. We thank Charisse Winston-Gray, PhD, and Floyd Sarsoza for their technical assistance with the MSD MULTI-SPOT Assay System. We thank Janna Wu for her technical assistance with microglial IHC quantification.

## AUTHOR CONTRIBUTIONS

D.S.W. contributed animal experimentation and data analysis. L.A. contributed tissue dissections, RNA isolation, qPCR and RNA-Seq analysis. H.R. contributed immunohistochemistry and data analysis. D.S.W., D.K.W., C.C. and P.D. contributed to experimental design. P.D. conceived and directed the study, contributed confocal microscopy, and supervised molecular work and data analysis. D.S.W., L.A., C.C., D.K.W. and P.D. contributed to manuscript preparation.

## MATERIALS & CORRESPONDENCE

Data requests and correspondence should be addressed to Dr. Paula Desplats

## SUPPLEMENTAL MATERIAL

### MATERIALS AND METHODS

#### Animals

APP23 transgenic (TG) mice (B6.Cg-Tg(Thy1-APP)3Somm/J; JAX Stock No: 030504) and non-transgenic (NTG) littermates control mice were housed in light-tight enclosures at the University of California, Los Angeles in the Laboratory of Circadian and Sleep Medicine. The mice were given *ad libitum* food (ALF; Teklad rodent diet 8604; Envigo, Indianapolis, IN) and water access until they were placed in the time-restricted feeding (TRF) or ad libitum feeding (ALF) cohorts. The work presented in this study followed all guidelines and regulations of the UCLA Division of Animal Medicine that are consistent with the Animal Welfare Policy Statements and the recommendations of the Panel on Euthanasia of the American Veterinary Medical Association.

#### Experimental design

Prior to any measurements or beginning any experiments, mice were habituated to a 12:12 light-dark (LD 12:12) cycle and single housing conditions in custom light-tight cabinets. We then measured diurnal rhythms in activity and sleep behavior from the mice. To assess circadian impairments in the mice, some animals were placed in constant darkness (DD), followed by light masking and 6 h phase advance protocols. Feeding manipulations were then performed on mice habituated to LD 12:12. Animals within each sex and genotype were randomly divided into two groups with one group continuing ALF and the other placed on TRF, with food available during 6 h in the middle of the mouse active phase, during zeitgeber time (ZT) 15 to ZT21. By definition, ZT0 is the time when lights go on and ZT12 is the time when the lights go off when the mice are in a 12:12 LD cycle. After habituating to the feeding protocol, metabolites were measured in the mice to ensure the treatment was inducing the expected metabolic effects. The mice remained on TRF or ALF food access until sacrificed for tissue collection. After all tests were completed and the mice had re-entrained to the LD cycle, mice within each genotype and treatment group were randomly assigned for sacrifice at one of four time points: ZT0, ZT6, ZT12, or ZT18. ZT0 and ZT6 tissues were collected in the light; ZT12 and ZT18 were collected in the dark. Brain tissue and blood serum were collected for analysis. This study used a total of 110 mice almost equally distributed across sex/ genotype/treatment throughout the study, and further grouped by ZT collection times. Treated mice started TRF at 6 to 9 months of age and were maintained on TRF until tissue collection, which occurred at 10 to 13 months of age.

#### Time-restricted feeding

Experimental mice were housed in standard cages with either normal ad libitum food or with a custom-made programmable food hopper that could temporally control access to food and prevent food consumption during restricted times. Feeders were open for 6 h in the middle of the mouse active phase (ZT15 to ZT21). Food consumption was initially measured twice per week to ensure that all mice acclimated to the new feeding regimen. Mice are coprophagic and were moved to new cages twice per week, as determined empirically, to diminish coprophagic behavior during the fasting period.

#### β-hydroxybutyrate and glucose measurements

Tail blood sampling was performed at ZT14 under dim red-light conditions (3 lux). Blood was tested for β-hydroxybutyrate (BOHB, 1.5 μL sample) and glucose (0.6 μL sample) using a commercially available glucose/ketone meter (Precision Xtra Blood Glucose and Ketone Monitoring System, Abbott Laboratories, Chicago, IL).

#### Monitoring of cage locomotor activity

Experimental mice were singly housed in cages with IR motion sensors (Honeywell, Charlotte, NC) and activity data were analyzed using the El Temps programs (A. Diez-Nogura, Barcelona, Spain; http://www.el-temps.com/principal.html) and ClockLab Analysis 6 (Actimetrics, Wilmette, IL). The data were recorded and analyzed as previously described ^71,72^. Mice were initially entrained to a 12:12 LD cycle. Locomotor activity was recorded using Mini Mitter (Bend, OR) data loggers in 3-min bins, and 7 to 10 days of data were averaged for analysis. Activity data were collected for a month prior to placement in total darkness or any feeding treatments. The amount of cage activity over a 24 h period was averaged and reported in arbitrary units (a.u.)/h. The number of activity bouts and the average length of bouts were determined, with a new activity bout defined after a gap of either 21 min (maximum gap: 21 min; threshold: 3 counts/min) or 1 min (maximum gap: 1 min; threshold: 3 counts/min). Circadian period length (Tau) was determined in DD, and re-entraining phase shifts were measured after a 6 h advance in the light/dark cycle (see below). Tau in DD was obtained from the slope of a line fitted to activity onset times. Phase shifts were calculated as: [(Δh “activity onset” to “entrained onset”)/days to entrainment] and presented as the shift in h/day.

#### Monitoring of immobility-defined sleep behavior

Immobility-defined sleep was determined as previously described ^71,72^. Mice were housed in standard see-through plastic cages containing bedding (without the addition of nesting material). A side-on view of each cage was obtained, with minimal occlusion by the food bin or water bottle, both of which were top-mounted. Cages were top lit using 850nm IR LED lights (LLH-850nm-60; LEDLightinghut, Shenzhen City, China). Video capture was accomplished using surveillance cameras with visible light filters (Gadspot Inc., City of Industry, CA) connected to a video-capture card (Adlink Technology Inc., Irvine, CA) on a custom-built computer system. ANY-maze software (Stoelting Co., Wood Dale, IL) was used for automated acquisition of mouse immobility.

Immobility was registered when 95% of the area of the animal remained immobile for more than 40 sec, which was previously determined to have 99% correlation with simultaneous EEG/EMG defined sleep ^73,74^. Continuous tracking of the mice was performed for a minimum of 5 sleep-wake cycles, with time-randomized visits (1 time/day) by the experimenter to confirm mouse health and video recording. Two sleep-wake cycles over 48 h were averaged for further analysis. Immobility-defined sleep data were exported in 1-min bins, and total sleep time was determined by summing the immobility durations in the rest phase (ZT0 to ZT12) and active phase (ZT12 to ZT24). An average waveform of hourly immobile-sleep over the two sleep-wake cycles was produced per sex, genotype, and treatment for graphical display.

#### Assessments of circadian function

To assess endogenous circadian period in the mice we measured free-running behavior in constant darkness for 10-14 days. To assess circadian response to external cues, mice were exposed to a single activity-augmenting dark pulse in the light phase (positive masking: 1 h dark pulse during the light phase at ZT3), then a single activity-suppressing light pulse in the dark phase (negative masking: 1 h light pulse in the dark phase at ZT15). Further, a 6 h phase advance was performed to assess circadian entrainment. Mice were permitted to recover for at least a week between light cycle manipulations, and entrainment was verified before each test was performed ^75^. These tests were not performed under TRF treatment, as TRF is itself a synchronizing stimulus (zeitgeber).

#### Tissue Collection

Mice were euthanized with isoflurane, either in the dark (ZT12 and ZT18) or in the light (ZT0 and ZT6). Brain hemispheres were collected and placed in either RNAlater Stabilization Solution (Thermo Fisher Scientific, Waltham, MA) or 4% paraformaldehyde. Serum was collected from blood incubated on ice for 30 min, centrifuged at 2000 g for 10 min at 4°C, and then stored until used at −80°C.

#### RNA isolation and RNA-seq analysis

Total RNA was isolated from hippocampus tissue dissected from one hemibrain per animal using RNeasy Lipid Tissue Mini kit (Qiagen, Hilden, Germany) as indicated by the manufacturer. Quality of the extracted RNA was assessed using TapeStation (Agilent Technologies, Inc., Santa Clara, CA). All samples showed RIN ≥8.

RNA-Seq library preparation was performed with poly-A enrichment using poly-T oligo-attached magnetic beads employing the PE150 sequencing strategy performed on a NovaSeq 6000 (Illumina, Inc., San Diego, CA) by Novogene Inc. (Sacramento, CA). Raw reads (>20M per sample) were imported to Galaxy platform for analysis ^76^. Reads were first cleaned by removing adapter sequences, trimming low-quality ends, and filtering reads with low quality using Trimmomatic. Sequence alignment of the resulting high-quality reads to the mouse reference genome (build GRCm38) was done using HISAT2 and quantification of gene-level expression was performed using featureCounts. DESEQ2 was used to generate normalized read counts and to determine differentially expressed genes. Genes with an average expression of more than 2 counts per minute (CPM) across all samples were considered as expressed. Pathway enrichment analysis was performed using Ingenuity Pathway Analysis (IPA) software (Qiagen), Metascape [http://metascape.org] ^77^, and Enrichr [https://maayanlab.cloud/Enrichr/] ^78^ web-based tools.

#### Quantification of mRNA by Real time PCR

Total RNA (1.0 μg) was used for reverse transcription to cDNA using a High-Capacity cDNA Reverse Transcription Kit (Applied Biosystems, Waltham, MA). Quantitative real-time PCR (Qpcr) was performed using TaqMan Fast Advanced Master Mix and mouse specific TaqMan probes (Thermo Fisher Scientific): *Arntl* (Mm00500226); *Per1* (Mm00501813); *Per2* (Mm00478113); *Cry1* (Mm00514392); *Sirt1* (Mm01168521); *Col1a2* (Mm00483888); *Dnajb4* (Mm00508908); *Kcnip4* (Mm00518835). qPCR reactions were performed in duplicate. Relative quantification of gene expression was calculated using β-actin (*Actb* (Mm00607939)) as an internal control and expressed as the inverse ratio to threshold cycle (1/dCt). AD-related genes were selected based on three sources: AMP-AD, DisGENET, and IPA AD-related molecules.

#### NanoString AD and Neuroinflammation Panels

Total RNA (1.0 μg) from hippocampus was used to hybridize either the nCounter^©^ Mouse AD Consortium Panel or the nCounter^©^ Mouse Neuroinflammation Panel (NanoString Technologies, Inc.). Raw data were exported into ROSALIND (version 3.35.10.0, https://rosalind.bio/) for analysis, including QC steps and differential gene expression. Normalization, fold changes and p-values were calculated using criteria provided by NanoString. ROSALIND® follows the nCounter® Advanced Analysis protocol of dividing counts within a lane by the geometric mean of the normalizer probes from the same lane. Housekeeping probes to be used for normalization are selected based on the geNorm algorithm as implemented in the NormqPCR R library (Perkins, JR. et al. BMC Genomics 13, 286 2012). Clustering of genes for the final heat-map of differentially expressed genes was done using the PAM (Partitioning Around Medoids) method using the fpc R library that takes into consideration the direction and type of all signals on a pathway, the position, role, and type of every gene, among other parameters (Hennig, C. Cran-package fpc. https://cran.r-project.org/web/packages/fpc/index.html).

#### Immunostaining

For immunohistochemistry (IHC), mice were sacrificed at the end of treatment and hemi-brains were extracted and fixed by 4% paraformaldehyde. Fixed brains were sectioned sagitally at 40 μm using a Leica VT1000S vibratome. Sections were washed three times in PBS, pre-treated with 1% Triton X-100, 10% H_2_O_2_ in PBS for 20 min at room temperature, washed again, and incubated for 1 h at room temperature in 10% serum according to secondary antibody species. The sections were incubated with primary antibodies to microglia marker Iba1 (1:500, FUJIFILM Wako Chemicals U.S.A. Corporation, code number 019-19741), NeuN (1:200, Milipore Corp., Cat. No. MAB377), 82E1 (1:500, Immuno-biological Laboratories, Cat. No. 10323), GFAP (1:200, Invitrogen 180063) at 4°C overnight. Sections were washed three times, incubated in 1:100 biotinylated secondary antibody (goat anti-rabbit, Vector Laboratories, BA-1000; or horse anti-mouse Vector Laboratories, BA-2000) for 30 min at room temperature, washed again, incubated in biotinylated HRP and avidin (ABC, Vector Laboratories, code number 30015 and 30016) for 1 h in the dark at room temperature and then treated with diaminobenzidine (DAB) Substrate Kit, Peroxidase (Vector Laboratories, SK-4100) for coloration. 20X images were collected using a Nanozoomer slide scanner (Humamatsu, Japan).

Quantification of each cell type was done in ImageJ using the “analyze particles” tool or determining the % area covered by the signal in hippocampus and cortex. Plaque count and area were manually acquired by blinded researchers using 82E1 DAB-stained sections exported into ImageJ.

#### Longitudinal labeling of plaques

To determine the rate of amyloid deposition, amyloid plaques were labelled *in vivo* on a subset of n=16 TG mice 1 mo after TRF protocol was started, by i.p. administration of Methoxy-X04 (Tocris Bioscience, Cat. No. 4920; 100 uL of 25 mg/mL in DMSO and diluted 1:10 in PBS) as previously reported ^47,79^. Fixed hemibrains collected after sacrificing the mice were later stained with anti-human amyloid beta antibody 82E1 (IBL 10323; 1:500 in PBST) using chicken anti-mouse Alexa Fluor 594. All Methoxy-X04 stained plaques were imaged on a Zeiss LSM 800 confocal microscope. Three-channel 12-bit images were sequentially acquired in 1 μM z-steps spanning the entire thickness of the plaque (Zeiss 63xplain apochromatic oil immersion lens, NA:1.4). Uniform pinhole AUM=1 was applied to all sections, with laser power ranging between 10 and 23% for the red (561 nm) and blue (405 nm) channels; digital gain ranging between 0.3 and 3.4 (red) and 0.3 and 2 (blue) and detector offset ranging for 1 to 2 in both channels. Settings were defined as to obtain the same exposure level based on pixel intensity in the blue channel (Methoxy-X04).

Plaques were manually quantified in ZEN Blue (Carl Zeiss AG) by blinded investigators who determined the areas detected in the red (82E1) and blue (Methoxy-X04) channels in orthogonal projection images generated for each plaque analyzed. A total of 16 APP23 TG mice and two sections per animal were analyzed (n=7 ALF; n=9 TRF; 0-7 plaque image/section/mouse).

#### Aβ40 and Aβ42 Quantification

Serum levels of Aβ40 and Aβ42 were determined in peripheral serum from n=38 mice using the Meso Scale V-PLEX Aβ Peptide Panel 1 (6E10) Kit (K15200E) processed on a MESO QuickPlex SQ 120 as indicated by the manufacturer, with final incubation at 4°C extended to overnight. NTG mice assayed (n=12-18) had no detectable levels of human Aβ40 or Aβ42 (not shown). Analysis was performed in the DISCOVERY WORKBENCH (Meso Scale Diagnostics, Rockville, MD).

#### Statistical analysis

Data were analyzed in GraphPad Prism 9 (GraphPad Software, Inc., San Diego, CA) unless indicated otherwise. Figure error bars represent the standard error of the mean (SEM) unless stated otherwise. For comparisons, t-tests were used to determine significant differences between means of two group values, either paired or unpaired as appropriate. To analyze sleep waveforms, we performed an analysis of variance (ANOVA) followed by multiple unpaired t-tests corrected for multiple comparisons using the Holm-Šídák method. Significance is indicated as follows: * = p≤0.05, ** = p≤0.01, *** = p≤0.001, **** = p≤0.0001. NanoString data for gene expression was analyzed using the Rosalind suite with P-value adjustment using the Benjamini-Hochberg method of estimating false discovery rates (FDR). Differential gene expression from RNAseq data was determined by DESEQ2 using the negative binomial distribution and a relaxed significance threshold of adj.p≤0.2 was used for discovery and pathway analysis. The periodicity of gene expression data was evaluated using the JTK_Cycle algorithm (Hughes et al., 2010) within the MetaCycle R package suite (Wu et al., 2016).

#### Data availability

RNA-seq data will be deposited at the NCBI Sequence Read Archive (SRA) and will be publicly accessible upon publication of the current manuscript.

